# Molecule template estimation using Validator Barcodes in multiplex PCR for adaptive immune repertoire profiling

**DOI:** 10.1101/2025.10.07.680989

**Authors:** Kitt D. Paraiso, Mark Izraelson, Alex Chenchik

## Abstract

TCR- and BCR-sequencing (TCR/BCR-seq) are two important technologies in studying the immune repertoire of samples such as PBMCs or tumors. In their most common form, these assays combine multiplex PCR of the repertoire using primers targeting regions of the V(D)J and the constant region with next-generation sequencing (NGS). The data produced by this assay provide a slew of information regarding immune repertoire(s) including the presence critical clonotypes, repertoire diversity, variable (V) gene usage, analysis of public clonotypes, etc. One issue that can arise during generation of the TCR/BCR-seq data is sequence bias during the PCR or NGS steps. To combat this, unique molecular identifiers (UMIs) have been used to identify and eliminate sequence bias. However, UMI fragments can be long and very diverse, resulting in the UMI sequences interfering with any of the multitude of primers during multiplex PCR. Here, we introduce Validator Barcodes (VBCs), a set of eight short barcodes (6-9 nucleotides in length). This compact set of barcodes improves PCR efficiency and facilitates PCR primer designs. Also, like UMIs, the VBCs may be used to estimate the number of template molecules (RNA or DNA). Using VBC-labeled primers for TCR and BCR repertoire profiling from PBMCs produces highly comparable results and similarly template values to those obtained through UMI-based assay counts. Overall, VBCs are a useful and simpler alternative to UMIs in assaying TCR and BCR repertoires.

## Introduction

The immune repertoire refers to the collection of B-cell receptors (BCRs) and T-cell receptors (TCRs) generated by the adaptive immune system to recognize a vast array of antigens. This repertoire is formed through the somatic recombination of gene segments encoding immunoglobulins and TCR chains, resulting in millions of unique clonotypes (Tonegawa, 1983; Roth, 2015). The diversity of the immune repertoire enables the immune system to detect and respond to numerous pathogens or cancer cells. Understanding the composition and dynamics of the immune repertoire is essential for insights into immune function, development, and disease.

Immune repertoire profiling or immunoprofiling involves high-throughput sequencing techniques to characterize the BCR and TCR repertoires at the genetic and/or transcriptomic levels (Seo and Choi, 2025). This technology allows the study of dynamics of clonotypes of interest or analysis at the repertoire level, which provides information on clonotype diversity, gene segment usage, etc. Applications of immunoprofiling include vaccine development, cancer immunotherapy, autoimmune disease research, infectious disease diagnostics, and transplantation immunology.

In certain cases, quantification of clonotypes rely on accurate counting of unique receptor molecules derived from individual templates. Unique molecular identifiers (UMIs) are random nucleotide tags incorporated during library preparation to label each original template molecule (König et al., 2010; Kivioja et al., 2012). UMIs help correct for PCR amplification bias and sequencing errors, thus providing more accurate quantification of clonotype abundance. Despite their advantages, UMIs present several challenges that can disrupt both experimental and bioinformatic procedures. UMI incorporation can introduce complexity in library preparation, increasing costs and technical variability. Errors in UMI sequences during PCR or sequencing can lead to incorrect collapsing of reads, resulting in incorrect template counts (Smith et al., 2017; Philpott et al., 2021; Ziegenhain et al., 2022; Sun et al., 2024). Incomplete UMI diversity can limit their effectiveness in highly complex repertoires (Clement et al., 2018). Additionally, in immunoprofiling studies where multiplex PCR is commonly used, the long and highly diverse sequences found in UMI can potentially interfere with amplification by the diverse primer pairs.

Here, we developed Validator barcodes (VBCs) as an alternative to UMIs, along with a normalization strategy to estimate the number of molecular templates. VBCs are short fragments from 6 – 8 nucleotides. These fragments are incorporated either during cDNA synthesis for RNA templates or barcode extension for DNA templates, both prior to PCR amplification. We find that VBC-based immunoprofiling yields improved PCR amplification compared to UMI-based immunoprofiling using the same set of gene-specific primers. VBC-based and UMI-based immunoprofiling generate similar repertoires from the same biological samples. We find that VBC-based assays yield similar quantification of molecular templates as UMI-based assays. Overall, our findings suggest that VBCs can act as an alternative to UMIs in immunoprofiling.

## Results

### Validator Barcodes for adaptive immunoprofiling

While UMIs have been used to improve accuracy in the quantification of clonotypes, authors have pointed out multiple issues with their usage including the length and randomness of their sequences (e.g. Clement et al., 2018; Ziegenhain et al., 2022). VBCs aim to address the later issues. Similar to UMIs, these sequences are incorporated during the initial steps of the molecular biology workflow (Figure 1A). For DNA templates, the VBCs are incorporated in the 3’ end of the amplicons (adjacent to the J region of the clonotypes) during a primer extension step. For RNA templates, the VBCs are incorporated in the 3’ end of the amplicons (adjacent to the C region of the clonotypes) during reverse transcription. In both cases, the result of the molecular workflow and next generation sequencing results in a paired end sequencing with read 2 containing VBCs.

**Figure 1.**
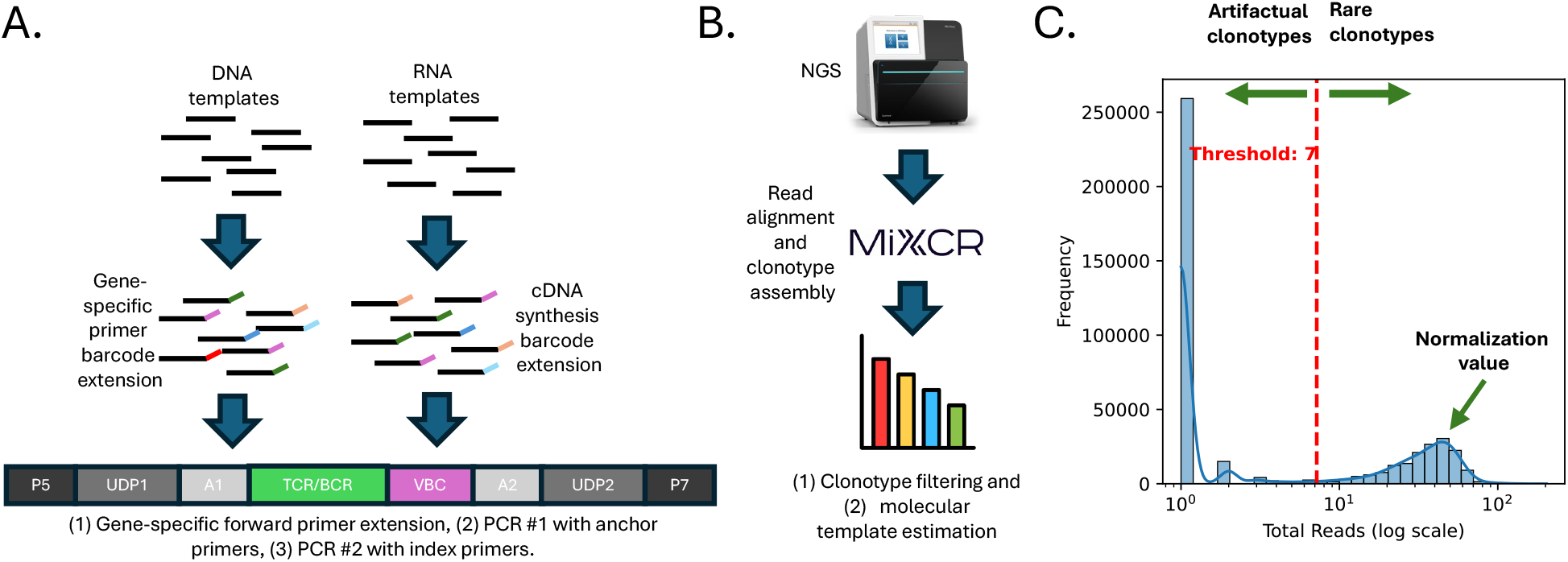
Immunoprofiling with Validator Barcodes. (A) Molecular workflow starts with the DNA and RNA templates. The validator barcodes (in color) are introduced either during gene-specific primer barcode extension in the DNA templates or during cDNA synthesis with gene-specific primers in the RNA templates. Both processes are followed by gene-specific forward primer extension and two PCRs. The resulting product for NGS contains the Illumina sequencing adapters (P5 and P7), Illumina dual index primers (UDP1 and UDP2), universal anchor primers (A1 and A2), the validator barcode, and the TCR or BCR amplicon. (B) bioinformatics workflow starts with the initial read alignment, followed by clonotype assembly using tools such as MiXCR. Clonotypes are filtered and passing clonotypes read counts are converted to estimated template count. (C) Distribution of reads per clonotype display a bimodal distribution where artifactual and real clonotypes are distinguished. For rare clonotypes appearing with only 1 type of validator barcode, the normalization value is identified. This value is then used as the factor to convert reads to estimated template counts.

The reads are then aligned to reference immune receptor genes and clonotypes are assembled (Figure 1B) with software such as MiXCR (Bolotin et al., 2015). Post-alignment, VBCs are used in two ways. First, the PCR and NGS steps can generate errors in the final sequences. VBCs can be used to identify sequences that are real vs artifactual. In this process, the assumption is that artifactual sequences have much lower abundance than real sequences. Second, VBCs can be used to identify a normalization factor to convert read counts to an estimate in the number of DNA or RNA molecule templates in the sample. During this process, it’s critical to identify clonotypes which only has one template (i.e. rare clonotypes). The read counts of this set of clonotypes create distribution (Figure 1C), the average of which is identified as the normalization factor. From there, the normalization is used as a denominator to convert the reads to the estimated number of templates.

With the reads aligned and clonotypes quantified, regular downstream analyses procedures for immunoprofiling data can be performed on the data. Repertoire statistics such as diversity metrics, gene usage, and clonotype overlaps can be measured. Clonotypes of interests such as peptide-activated clonotypes, vaccine-associated clonotypes, sequences of tumor infiltrating lymphocytes, etc. could be identified. The immunoinformatics community have created a slew of tools to perform these analyses (Mhanna et al., 2024).

### Clonotypes in the VBC and UMI assays

To test out the performance of VBCs, we’ve performed immunoprofiling for (A) TCR and BCR, (B) RNA and DNA template based assays, (C) CDR3 amplification and full length amplification and (D) human and mouse PBMCs (see Table 1 for full list of samples). In addition, we generated four samples using UMIs for comparison. These four UMI samples matches one of the VBC samples in exact conditions including biological sample used, amount of nucleic acid material, and molecular workflow. During the PCR amplification steps, we found that the VBC showed improved efficiency (Figure 2A) where we see stronger bands, in particular for the DNA assay. This could likely be attributed to the shorter sequences in the VBCs vs UMIs.

**Table 1.** Experiment sample list. List of samples in the experiment. Sample names are formatted as Stemplate $ceUType-$amplicon-$version-$templateAmount. Template is either R for RNA or D for DNA. Mouse cell types (mT and mB) are distinguished from human cell types (T and B) with an’m’ before the cell type. Amplicon is either CDR3 for CDR3 or CDR123 for full length. Version is either VBC orUMI.

| Samples | Version | Species | Amplicon | Template | Template Amount |
| --- | --- | --- | --- | --- | --- |
| R_T-CDR123-VBC-50ng | VBC | human | Full Length | RNA | 50 ng |
| R_T-CDR123-VBC-10ng | VBC | human | Full Length | RNA | 10 ng |
| R_T-CDR3-VBC-50ng | VBC | human | CDR3 | RNA | 50 ng |
| R_T-CDR3-VBC-10ng | VBC | human | CDR3 | RNA | 10 ng |
| R_B-CDR123-VBC-50ng | VBC | human | Full Length | RNA | 50 ng |
| R_B-CDR123-VBC-10ng | VBC | human | Full Length | RNA | 10 ng |
| R_B-CDR3-VBC-50ng | VBC | human | CDR3 | RNA | 50 ng |
| R_B-CDR3-VBC-10ng | VBC | human | CDR3 | RNA | 10 ng |
| R_mT-CDR3-VBC-10ng | VBC | mouse | CDR3 | RNA | 10 ng |
| R_mB-CDR3-VBC-10ng | VBC | mouse | CDR3 | RNA | 10 ng |
| D_T-CDR3-VBC-2ug | VBC | human | CDR3 | DNA | 2 µg |
| D_T-CDR3-VBC-0p4ug | VBC | human | CDR3 | DNA | 0.4 µg |
| D_B-CDR3-VBC-2ug | VBC | human | CDR3 | DNA | 2 µg |
| D_B-CDR3-VBC-0p4ug | VBC | human | CDR3 | DNA | 0.4 µg |
| D_B-CDR3-UMI-2ug | UMI | human | CDR3 | DNA | 2 µg |
| D_T-CDR3-UMI-2ug | UMI | human | CDR3 | DNA | 2 µg |
| R_B-CDR3-UMI-50ng | UMI | human | CDR3 | RNA | 50 ng |
| R_T-CDR3-UMI-50ng | UMI | human | CDR3 | RNA | 50 ng |

**Figure 2.**
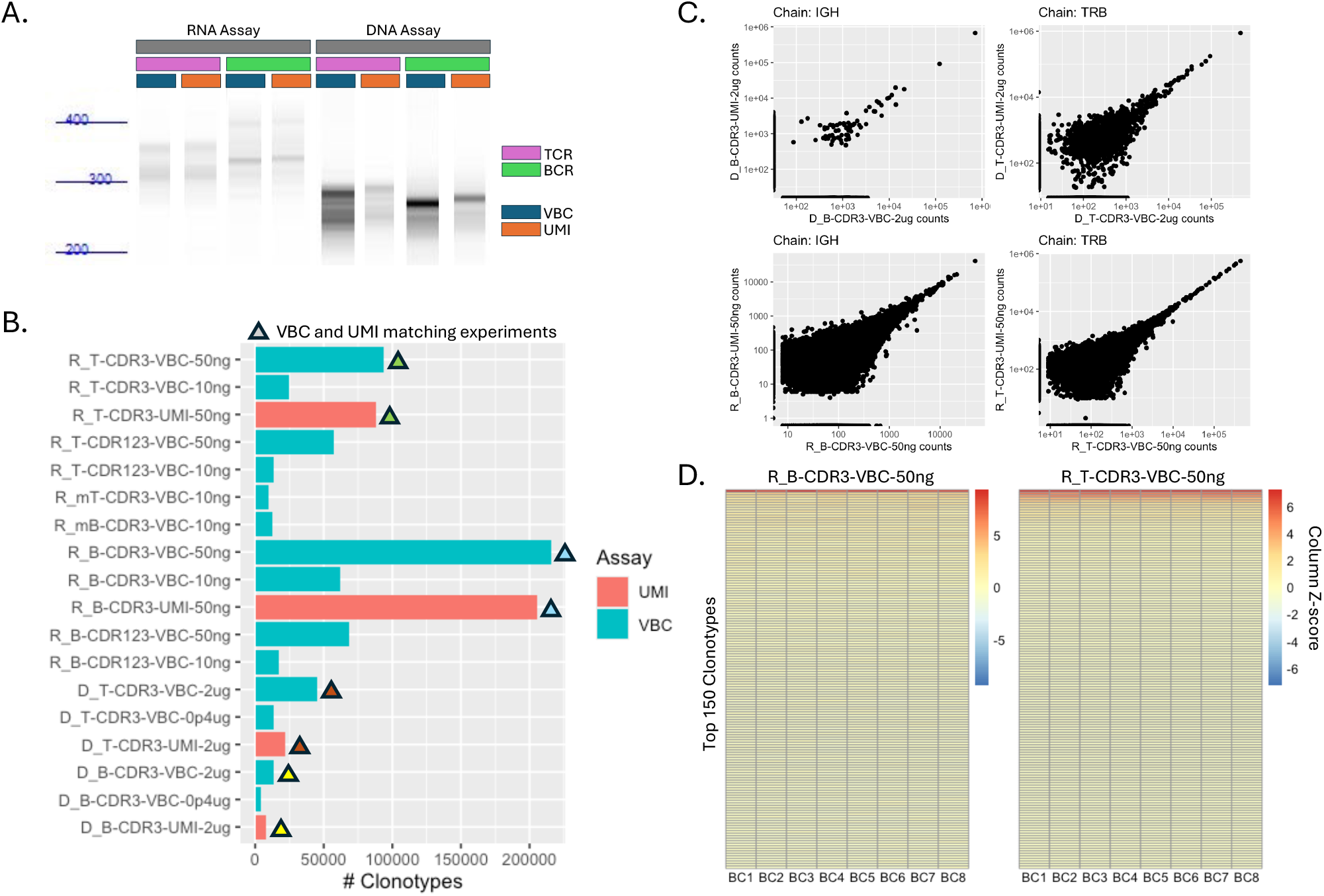
Repertoire from VBC and UMI based assays. (A) PCR amplification of VBC- and UMI-based samples. (B) Number of clonotype identified. (C) Reproducibility between VBC and UMI assays in IGH and TRB chains. (D) Heatmap of read counts across VBCs from RNA templates.

These samples are analyzed using standard MiXCR presets (see Methods). Both VBC and UMI samples yielded great alignment rates at 90+% for samples with CDR3 amplicons. Unsurprisingly, the full-length amplicon samples yielded lower alignment rates since they required longer sequencing (600 NT PE vs 300 NT PE). Since the barcode sequences present in the VBC experiments were pre-selected, we were concerned about the performance of each barcode during amplification relative to each other. However, we found no evidence that any of the barcodes showed any bias as the reads in our samples showed similar distribution across 8 barcodes (Figure S1).

The repertoire in the UMI and the VBC assays were generally similar. The number of clonotypes in corresponding VBC and UMI samples were similar in number (Figure 2B). The read counts were similar between the UMI and VBC samples as well across multiple receptor chains IGH and TRB (Figure 2C); and IGK, IGL and TRA (Figure S2). Read counts correlated well between the repertoires of corresponding samples. This is especially true for the clonotypes of high abundance.

Since the VBCs categorize reads into one of the unique sequences, the setup effectively simulates 8 technical replicates during the molecular workflow. The octuplicates can be used to assess variation in clonotype quantification. At least for the top clonotypes, the read quantitation reproduces well between the simulated technical replicates (Figure 2D, Figure S3). This is more apparent in the RNA samples where there are more clonotypes observed.

### Repertoire metrics in VBC and UMI assays

Analyzing both TCR and BCR repertoire metrics is fundamental for understanding immune system dynamics and responses. Changes in the gene usage (i.e. the frequency of V, D, and J gene segments) could be indicative of pathogenic diseases, autoimmune diseases or cancer. For example, particular gene segments have been associated with systemic lupus erythematosus (Ota et al., 2023), COVID-19 (Schultheiß et al., 2020), chronic hepatitis B infection (Qu et al., 2016), lymphocytic choriomeningitis virus infection (Kuhn et al., 2022), murine cytomegalovirus infection (Kuhn et al., 2022) and rheumatoid arthritis (Turcinov et al., 2023). Similarly, the diversity of TCR and BCR repertoires provide insights into how the immune system adapts to these challenges such as that seen in malaria infection (Frimpong et al., 2022), acute respiratory symptoms in infants (Paschold et al., 2023), aging (Yager et al., 2008) and the immune system’s ability to fight CMV infection (Wang et al., 2012).

In our experiments, the V gene segment usage is similar between the UMI and VBC assays of the same biological sample (Figure 3A). A handful of V genes are robustly present in each sample, while other genes are less so. Clonotype occupancy is similar between the UMI and VBC samples, as well (Figure 3B). The DNA samples tend to have the top clonotypes occupy more of the repertoire than the RNA samples. This is consistent with the D50 diversity metric, which calculates the number of clonotypes that occupy the top 50% of the repertoire space (Figure 3C). The true diversity metric calculates diversity by counting the number of clonotypes along with their relative abundances to account for both the richness and evenness of clonotypes in the repertoire. This metric also shows consistency between the UMI and VBC samples (Figure 3D). Overall, the UMI and VBC assays recapitulate each other’s results at the clonotype-level and at the repertoire-level.

**Figure 3.**
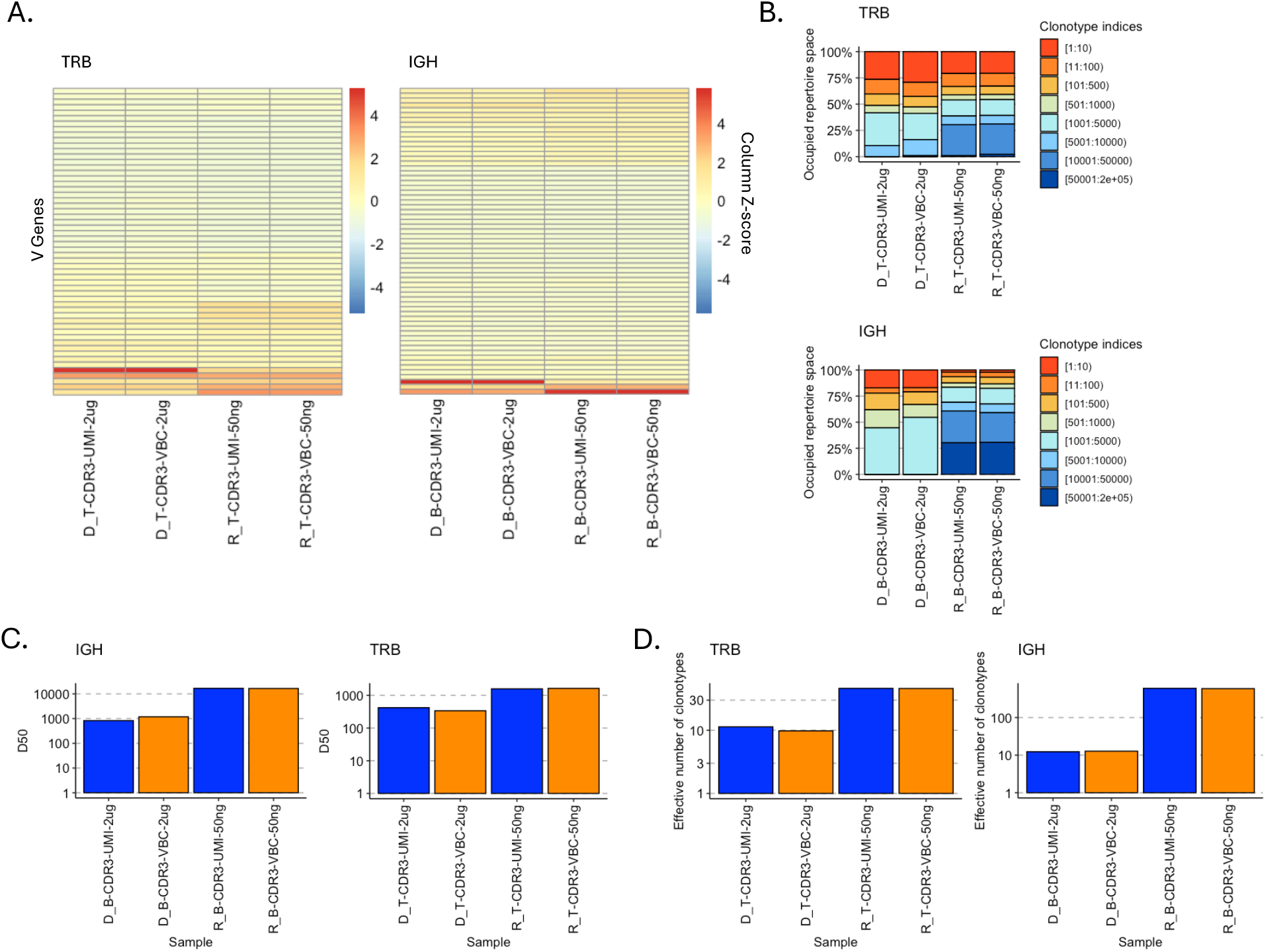
Repertoire metrics. (A) Z-score normalized read counts of V genes in the repertoire for TRB and IGH. (B) Clonotype occupancy of the repertoire divided by ranked read counts (e.g. top 1 – 10, top 11 – 100). (C) Diversity metric measured with the D50 index. (D) Diversity metric measured with the True diversity index.

### Template estimation using VBCs

The number of templates can be acquired from the UMI-based assay through counting of unique UMI sequences. For the VBC-based assay, the number of templates is estimated. For this procedure, we first identify clonotypes that are only found in a single VBC bin. Statistically, these clonotypes are rare and most likely, the reads from these clonotypes originated from a single template. After filtering out clonotypes the sequences that are artifactual, we identify the average of the distribution of the reads originating from a single template (Figure 4A). This value is the number of reads per one template. This is then used as the denominator to convert read counts to estimated number of DNA or RNA templates.

**Figure 4.**
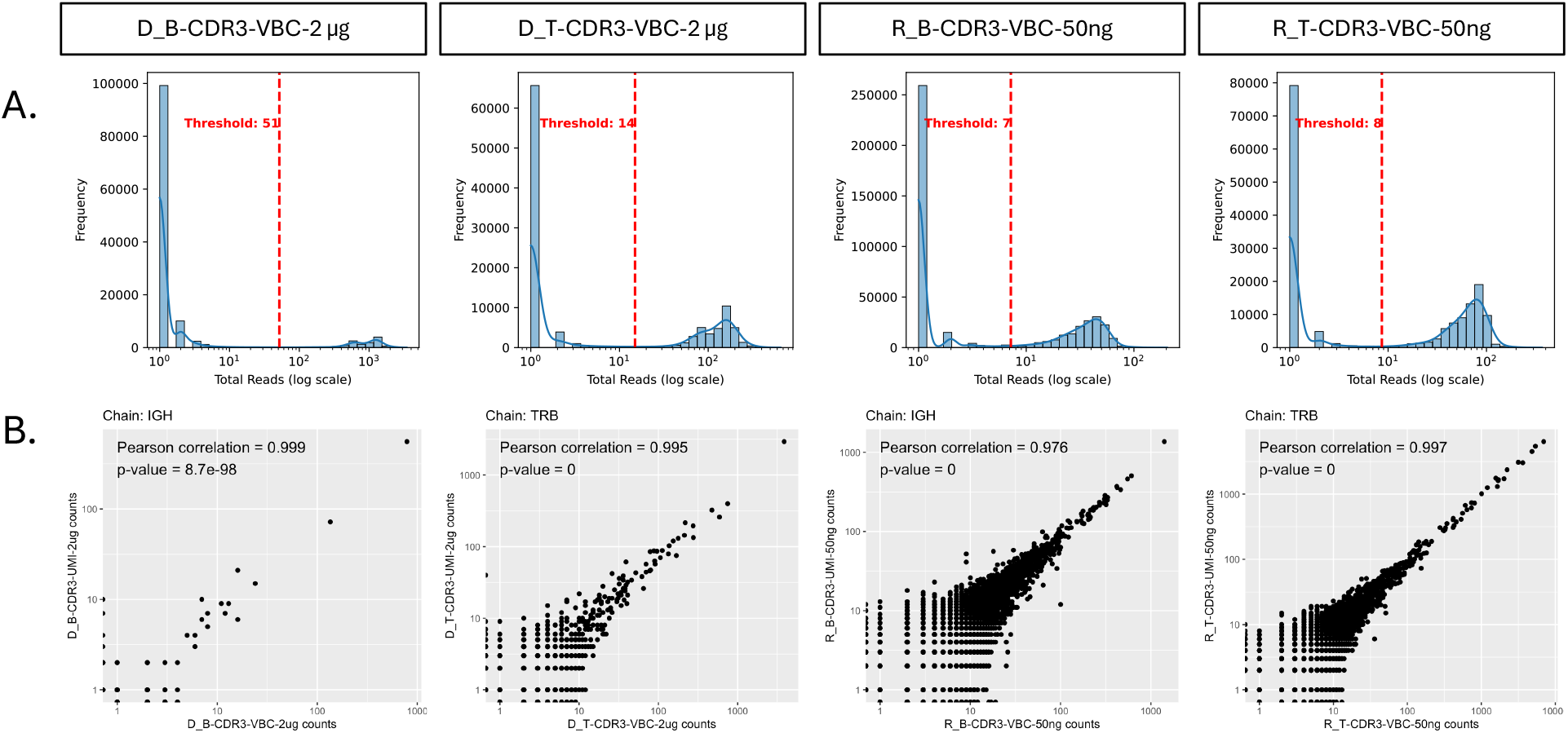
Template estimation using VBCs. (A)Histograms of the number of read counts for each clonotype that was only found in 1 VBC. The threshold value identifies the cutoff to distinguish real vs artifactual clonotypes. The average value of the left peak is used as the normalization value to convert read counts to UMIs. (B) Scatterplot of the UMI count (in the UMI experiments) vs the estimated number of templates (in the VBC experiments).

We compared the values estimated from the VBC-based assay to the UMI-based assay for the cases where a corresponding sample exists (Figure 4B). For the four cases that we compared, we found that the number of UMI and the estimated number of templates correspond extremely well, especially in the counts of 100+ range. We still see great correlation in the 10 - 100 counts range, while at the < 10 range, we start to see more variation between the two assays. This variation at lower abundant clonotypes is unsurprising as it can be attributed to less robust sampling of template material when preparing the sample for the pre-NGS molecular workflow. Additionally, it could be due to dropout events during template capture in the cDNA synthesis in the RNA-based assay or the primer elongation step in the DNA-based assay. This data suggests that for practical purposes, the VBCs are subicient to determine the number of RNA and DNA templates during immunoprofiling.

## Discussion

Immune repertoire profiling has become indispensable in biomedical research, providing insights into adaptive immune responses across diverse conditions. By analyzing TCR and BCR repertoires, researchers have uncovered clonal patterns in cancer, autoimmune disorders, and infectious diseases. This in turn enables the development of targeted diagnostics and therapies. Here, we have developed Validator Barcodes (VBCs) as an alternative to UMIs. VBCs are low diversity, short nucleotide fragments that can be used to estimate the number of molecules in the data. We find that the use of VBCs generate similar clonotypes, repertoire metrics, and molecular template quantification as UMIs with the same set of TCR and BCR gene segment primers. In addition, we find that the shorter VBC fragments allow for better PCR amplification of clonotypes.

## Supplemental Figures

**Figure S1.**
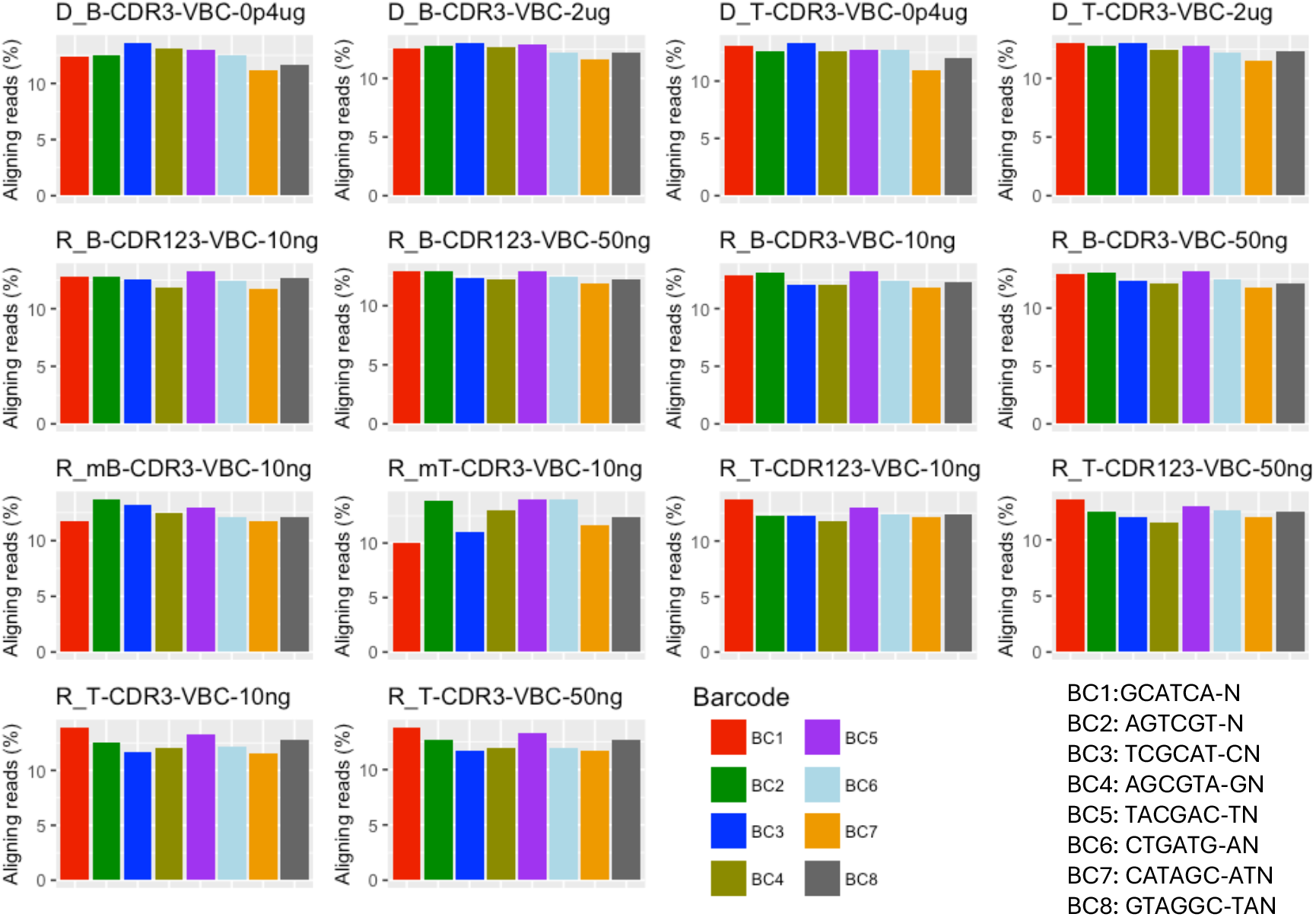
Read distribution across VBCs.

**Figure S2.**
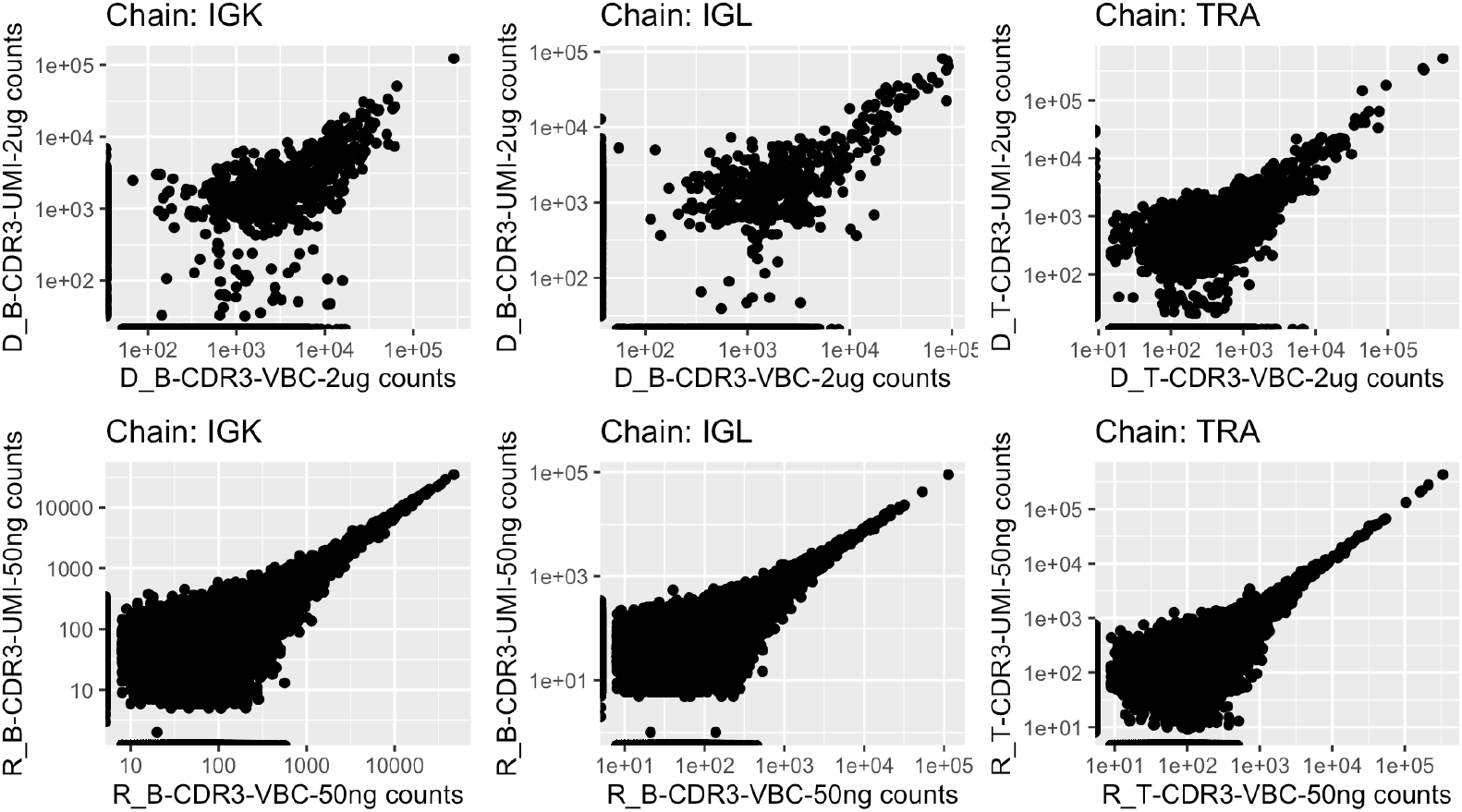
Reproducibility between VBC and UMI assays in IGK, IGL and TRA chains.

**Figure S3.**
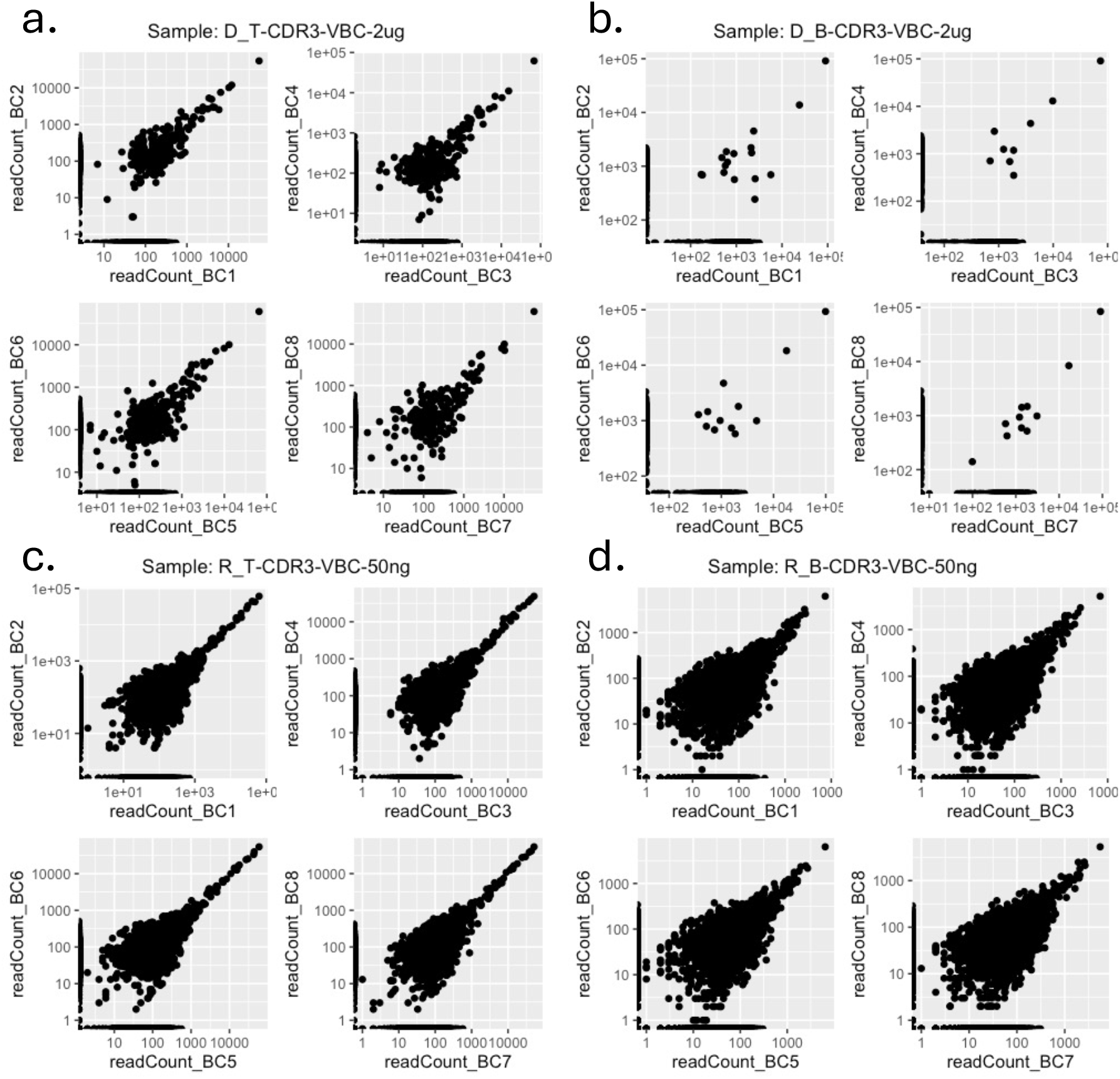
Reproducibility between VBC barcodes in IGH and TRB chains for B cell and T cell samples, respectively.

## Data Availability

The datasets generated for this study are available under accession number GSE297422. The MiXCR preset for analysis of the VBC-based assays are available here. Additional data and analyses supporting the findings of this study are available from the corresponding author upon request.

## Methods

### Adaptive immune response profiling

#### Nucleotide extraction

QIAGEN RNAeasy Micro/Mini Plus Kit for total RNA isolation. DNeasy Blood & Tissue kit was used for DNA extraction.

#### DNA- and RNA-based immunoprofiling

Cellecta DriverMap Adaptive Immune Receptor (AIR) was used to perform TCR and BCR repertoire profiling.

### Bioinformatic analysis

#### Sample Demultiplexing

*bcl2fastq* (Illumina, Inc., 2019) was used to demultiplex and convert the sequencing results to individual sample fastqs.

#### QC Analysis

*fastqc* (Andrew, 2010) was used to assess the quality of the sequencing reads prior to downstream analysis.

#### Read Alignment

*mixcr* (Bolotion et al., 2015) was used to align the reads to the reference immune repertoire genes. In addition, this command also generates the clonotype tables along with relevant metadata information.

The command used for the VBC-based assays is:

~~~
mixcr analyze local:cellecta-new-bulk-kit \
  --species $SPECIES \
  -–assemble-clonotypes-by $ASSEMBLY_TYPE \
  $SPLIT_CLONES_CONFIG \
  --$MOLECULE \
  --tag-pattern $TAG_PATTERN \
  --rigid-left-alignment-boundary \
  --floating-right-alignment-boundary $BOUNDARY \
  --export-productive-clones-only \
  $READ_1 $READ_2 $OUTPUT_NAME
~~~

For UMI-based assays, the command used is:

~~~
mixcr analyze $PRESET \
  --species $SPECIES \
  --export-productive-clones-only \
  $READ_1 $READ_2 $OUTPUT_NAME
~~~

where

`$SPECIES = hsa` (human) or `mmu` (mouse)

`$ASSEMBLY_TYPE = CDR3` (CDR3 amplicon) or `{CDR1Begin:FR4End}` (Full length amplicon)

`$SPLIT_CLONES_CONFIG = --split-clones-by C` (B cells) or leave blank (T cells)

`$MOLECULE = rna` (for RNA) or `dna` (for DNA)

`$BOUNDARY = J` (CDR3 amplicon) or `C` (Full length amplicon)

`$READ_1, $READ_2` = paired end read file names

`$OUTPUT_NAME` = name of output files

`$PRESET = cellecta-human-rna-xcr-umi-drivermap-air` (RNA-based) or `cellecta-human-dna-xcr-umi-drivermap-air` (DNA-based)

The `$TAG_PATTERN` is set as:

~~~
‘^(R1:^*^)\^(MIVBC:GCATCAN)(R2:^*^)|^(MIVBC:AGTCGTN)(R2:^*^)|^(MIVBC:TCGCATCN)(R2:^*^)|^(MIVBC:AGCGTAGN)(R2:^*^)|^(MIVBC:TACGACTN)(R2:^*^)|^(MIVBC:CTGATGAN)(R2:^*^)|^(M IVBC:CATAGCATN)(R2:^*^)|^(MIVBC:GTAGGCTAN)(R2:^*^)’
~~~

The preset for the VBC-based assay is specified by a cellecta-new-bulk-kit.yaml file.

#### Template Estimation

The number of molecule templates is estimated using a normalization factor. Briefly, rare clonotypes or clonotypes originating from a single template are identified from each dataset. First, the clonotypes are binned by as to how many VBCs they are identified with (from 1 – 8 VBCs). The clonotypes that were only identified in a single VBC (nVBC = 1) are further processed. The read counts from this group are plotted whereby two distributions are identified. The first distribution is an exponential decay originating from zero, and the second distribution is a normal distribution at some positive value. To distinguish the real clonotypes (second distribution) and the artifactual clonotypes (first distribution), kernel density estimation is used to smoothen the distribution. Then the two maximas are identified corresponding to the two distributions. Right in between the maximas is the lowest point, which is used as a filtering cutoff to eliminate the artifactual clonotypes. The filtered set of nVBC = 1 are then determined as the set of rare clonotypes. The second peak, which corresponds to the average of the filtered nVBC = 1, is used as the normalization factor to determine the estimated number of templates. This procedure is performed all types of VBC-based immunoprofiling assays in this study. The code to estimate the number of templates is available here.

#### Downstream Analysis

The *immunarch* (ImmunoMind Team, 2019) package is used for downstream analysis to study gene usage, repertoire diversity and clonotype occupancy. R (R Core Team, 2023) is used for various statistical analyses and data visualization.

